# A Multi-State Model of the CaMKII Dodecamer Suggests a Role for Calmodulin in Maintenance of Autophosphorylation

**DOI:** 10.1101/575712

**Authors:** Matthew C. Pharris, Thomas M. Bartol, Terrence J. Sejnowski, Mary B. Kennedy, Melanie I. Stefan, Tamara L. Kinzer-Ursem

## Abstract

Ca^2+^/calmodulin-dependent protein kinase II (CaMKII) accounts for up to 2 percent of all brain protein and is essential to memory function. CaMKII activity is known to regulate dynamic shifts in the size and signaling strength of neuronal connections, a process known as synaptic plasticity. Increasingly, computational models are used to explore synaptic plasticity and the mechanisms regulating CaMKII activity. Conventional modeling approaches may exclude biophysical detail due to the impractical number of state combinations that arise when explicitly monitoring the conformational changes, ligand binding, and phosphorylation events that occur on each of the CaMKII holoenzyme’s twelve subunits. To manage the combinatorial explosion without necessitating bias or loss in biological accuracy, we use a specialized syntax in the software MCell to create a rule-based model of the twelve-subunit CaMKII holoenzyme. Here we validate the rule-based model against previous measures of CaMKII activity and investigate molecular mechanisms of CaMKII regulation. Specifically, we explore how Ca^2+^/CaM-binding may both stabilize CaMKII subunit activation and regulate maintenance of CaMKII autophosphorylation. Noting that Ca^2+^/CaM and protein phosphatases bind CaMKII at nearby or overlapping sites, we compare model scenarios in which Ca^2+^/CaM and protein phosphatase do or do not structurally exclude each other’s binding to CaMKII. Our results suggest a functional mechanism for the so-called “CaM trapping” phenomenon, such that Ca^2+^/CaM structurally excludes phosphatase binding and thereby prolongs CaMKII autophosphorylation. We conclude that structural protection of autophosphorylated CaMKII by Ca^2+^/CaM may be an important mechanism for regulation of synaptic plasticity.

**Author summary:** In the hippocampus, the dynamic fluctuation in size and strength of neuronal connections is thought to underlie learning and memory processes. These fluctuations, called synaptic plasticity, are in-part regulated by the protein calcium/calmodulin-dependent kinase II (CaMKII). During synaptic plasticity, CaMKII becomes activated in the presence of calcium ions (Ca^2+^) and calmodulin (CaM), allowing it to interact enzymatically with downstream binding partners. Interestingly, activated CaMKII can phosphorylate itself, resulting in state changes that allow CaMKII to be functionally active independent of Ca^2+^/CaM. Phosphorylation of CaMKII at Thr-286/287 has been shown to be a critical component of learning and memory. To explore the molecular mechanisms that regulate the activity of CaMKII holoenzymes, we use a rule-based approach that reduces computational complexity normally associated with representing the wide variety of functional states that a CaMKII holoenzyme can adopt. Using this approach we observe regulatory mechanisms that might be obscured by reductive approaches. Our results newly suggest that CaMKII phosphorylation at Thr-286/287 is stabilized by a mechanism in which CaM structurally excludes phosphatase binding at that site.

## Introduction

CaMKII is a protein of interest because of its crucial role in synaptic plasticity [1–5]. In the hippocampus, synaptic plasticity in the post-synapse occurs within mushroom-shaped protrusions called dendritic spines [6]. Synaptic plasticity is dependent on calcium ion (Ca^2+^) flux through N-methyl-D-aspartate receptors (NMDARs) located on the dendritic spines of the post-synaptic neuron [7]. Depending on the magnitude, frequency, and location of Ca^2+^ flux, synaptic plasticity may produce increases or decreases (or neither) in synaptic strength [8, 9]. Large, higher-frequency Ca^2+^ spikes can induce an enduring up-regulation of synaptic strength, called long-term potentiation (LTP); while weak, lower-frequency Ca^2+^ spikes can induce an enduring down-regulation of synaptic strength, called long-term depression (LTD) [9, 10]. Whether Ca^2+^ spikes induce LTP or LTD depends on relative activation of intracellular protein signaling networks. When Ca^2+^ first enters the dendritic spine, it interacts with a variety of buffer and sensor proteins, chiefly calmodulin (CaM), which has many protein targets in the spine, including CaMKII [5, 11, 12].

The CaMKII holoenzyme contains at least twelve subunits [13–16] arranged as two rings of six. As shown in Fig 1, each CaMKII subunit features an N-terminal kinase domain and C-terminal hub domain [17]. Between the kinase and hub domains is a flexible regulatory domain which lends to the subunit a wide range of movement away from the holoenzyme’s central hub. A crystal structure of human alpha-CaMKII expressed in *E. coli* published by Chao *et al*. (2011) shows CaMKII subunits as able to rapidly and stochastically pivot between a “docked” and “undocked” conformation, seemingly mediated by residues on the kinase domain’s activation loop and a spur structure on the hub domain (see Fig 3C in [17]), such that a docked subunit may be inaccessible to CaM binding. In contrast, a more recent work using electron microscopy with rat alpha-CaMKII expressed in Sf9 cells suggests that less than 3 percent of subunits exhibit a compact (or docked) conformation [18]. Given the uncertainty in the field, we include subunit docking and undocking in our model, allowing for future exploration of this possible subunit functionality. In addition to docking and undocking, each subunit can be in an “inactive” conformation when the regulatory domain is bound to the kinase domain (Fig 1B), or an “active” conformation when this binding is disrupted by the binding of Ca^2+^/CaM or phosphorylation at Thr-286 [17, 19]. In the active conformation the catalytic domain of a subunit is able to bind and phosphorylate enzymatic substrates. A subunit may spontaneously return to an inactive conformation in the absence of Ca^2+^/CaM or phosphorylation at Thr-286 [19].

**Fig 1.**
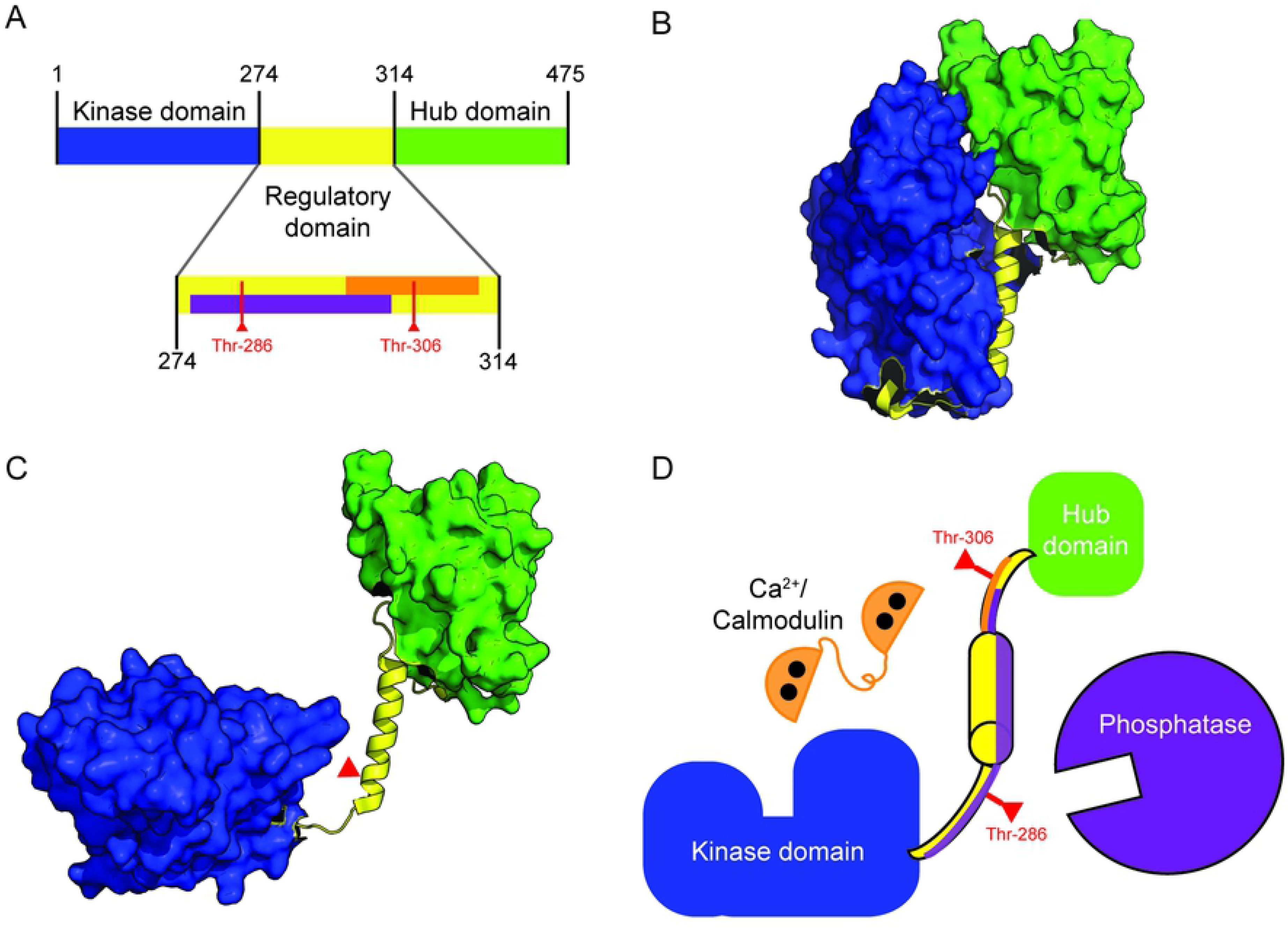
Schematic of CaMKII Subunit Structure. (A) Map of amino acid residues in a CaMKII subunit. The N-terminal kinase domain (blue) approximately spans residues 1-274. The regulatory domain (residues 275-314, yellow) binds to the kinase domain autoinhibiting the kinase activity of the each CaMKII subunit. The putative phosphatase binding site is also shown purple. The Ca^2+^/CaM binding site is shown in orange. Subunits self-associate via the hub domain (residues 315-475, green) to form multimeric complexes of 12-14 subunit holoenzymes. (B) The “inactive” CaMKII subunit (PDB: 3SOA) in which the regulatory domain (yellow) is closely associated with the kinase domain (blue). (C) A schematic of the “active” CaMKII subunit. The regulatory domain (yellow) is not bound to the kinase domain (blue). This schematic was generated by manually modifying PDB entry 3SOA to illustrate how the regulatory domain may be available for Ca^2+^/CaM binding and the kinase domain open for substrate binding. (D) Cartoon depiction of all protein species in our model, in which Ca^2+^/CaM (orange) or phosphatase (purple) may bind to the regulatory domain (yellow) of a CaMKII subunit.

CaMKII activity can become Ca^2+^/CaM-independent through phosphorylation at Thr-286, which is required for LTP [3, 20]. Importantly, this phenomenon is an autophosphorylation: it is thought to occur when an active subunit phosphorylates neighboring subunits within the same holoenzyme [21, 22]. Autophosphorylation at Thr-286 (“pThr-286”) is thought to provide structural stability to a subunit’s active conformation (reviewed in [23]) [24]. Because CaMKII plays a key role in the induction of LTP, and ultimately learning and memory (reviewed in [4, 8]), we seek to better understand the biochemical regulation of CaMKII activation and autophosphorylation via computational modeling.

To characterize the spatiotemporal regulation of CaMKII, experimental studies are increasingly complemented by computational models [15, 17, 25, 26]. Computational models of Ca^2+^-dependent signaling implicate competition, binding kinetics, feedback loops, and spatial effects in regulating enzyme activation [7, 12, 24, 27, 28]. However, fully characterizing these and other mechanisms of CaMKII regulation is impeded by the challenge of accurately portraying the CaMKII holoenzyme. As described by previous work, combinatorial explosion applies to models of CaMKII (and similar biomolecules) because the protein exhibits a large number of functionally significant and not necessarily inter-dependent states [24, 26, 29–31]. The large number of possible states of CaMKII can neither be explicitly specified nor efficiently evaluated with conventional mass action-based methods. Indeed, for just one CaMKII hexamer ring, we estimate a state space of ∼32 billion states, and for the full dodecamer approximately 10^20^ possible states (See S1 Appendix). The numbers of possible CaMKII states far exceeds the number of CaMKII molecules in a dendritic spine, suggesting that some states never occur and are therefore not functionally important. Previous models leverage this observation to reduce the model state space and provide valuable insight to CaMKII binding and autophosphorylation dynamics [24, 31–34]. However, for CaMKII it remains unclear which states functionally participate in synaptic plasticity. Reduced models can inadvertently obscure key mechanisms regulating CaMKII activation and autophosphorylation. To elucidate complex regulatory mechanisms, it may be necessary for models to provide for all possible states *ab initio*.

In this work, we use rule-based model specification and particle-based rule evaluation methods to overcome combinatorial explosion [26, 30, 35]. Rules are conditions, based primarily on experimental observations, that prescribe when an implicitly-defined reaction may occur. At a given iteration, only states that matter for the execution of a particular rule are explicitly declared. States that do not matter to a particular rule can be omitted, a principle that has been paraphrased as “don’t care, don’t write” [36]. We use rule- and particle-based methods within the spatial-stochastic software MCell 3.3 [28, 37] to present a comprehensive multi-state model of the CaMKII dodecamer. Other simulation platforms can also overcome combinatorial explosion through rule-based model specification (e.g. BioNetGen [38]) or network-free approaches (e.g. NFsim [39]). Unlike other platforms, MCell 3.3 provides both spatial-stochastic and rule-based modeling, although multi-state molecules in MCell 3.3 cannot diffuse. We use MCell 3.3 in anticipation of future MCell versions accounting for multi-state molecule diffusion, and to eventually build on simulations with physiological dendritic spine geometries such as those by Bartol *et al*. (2015) [40].

Here, we validate this rule-based MCell model of CaMKII regulation against current descriptions of the Ca^2+^ frequency-dependence of CaMKII activation. By varying the rules and model parameter values we can simulate different experimental manipulations of CaMKII interaction with Ca^2+^/CaM and phosphatase and thereby explore various mechanisms regulating CaMKII activity. In particular, we show that Ca^2+^/CaM is important not only for regulating activation of CaMKII but may also contribute to the maintenance of CaMKII phosphorylation at Thr-286. We hypothesize that by limiting access of phosphatases to CaMKII Thr-286 (perhaps by steric hindrance), Ca^2+^/CaM may prolong the lifetime of the auto-phosphorylated state.

## Results

### Model Development

#### Molecular Species

The model contains three protein species: CaM, protein phosphatase, and CaMKII. Ca^2+^/CaM facilitates CaMKII activation, which leads to autophosphorylation at Thr-286, and phosphatase activity facilitates de-phosphorylation at Thr-286. Both protein phosphatase 1 (PP1) and protein phosphatase 2A (PP2A) have been shown to dephosphorylate Thr-286, though in different subcellular fractions (reviewed by [21, 41–43]). Here we refer to them generally as protein phosphatase (PP).

CaM and PP are modeled in MCell as conventional cytosolic molecules. CaM is modeled as having one of two states: un-bound apo-CaM and fully-saturated Ca^2+^/CaM (four Ca^2+^ bound to CaM). Although we and others have described the importance of sub-saturated Ca^2+^/CaM states with fewer than 4 Ca^2+^ [12, 24, 31, 44–46], the dynamics of Ca^2+^-CaM binding and the binding of sub-saturated Ca^2+^/CaM to CaMKII are beyond the scope of this current work. Indeed, accounting for sub-saturated Ca^2+^/CaM would here require a multi-state representation, and because multi-state molecules cannot diffuse in MCell 3.3, we simplify our Ca^2+^/CaM model to allow CaM and CaMKII to interact. Thus, similarly to previous models [27, 47], we assume that apo-CaM has a negligible affinity for CaMKII; only fully-saturated Ca^2+^/CaM binds CaMKII. In contrast to CaM, PP is modeled as single-state protein that is constitutively active and able to bind auto-phosphorylated CaMKII subunits. Our representation of constitutively active PP is consistent with previous models such as that by Lisman and Zhabotinsky (2001) [48].

CaMKII is modeled as a multi-subunit complex, defined using a specialized model syntax for complex molecules (COMPLEX_MOLECULE) in MCell 3.3 [49]. This syntax allows for explicit representation of individual CaMKII dodecamers with distinguishable subunits. As shown in Fig 2, the holoenzyme is arranged as two directly-apposed, radially-symmetric rings each with six subunits. Each subunit features five “flags”, each standing for a particular state that each CaMKII subunit can adopt. Flags are used in rule evaluation, which occurs at each time step and for each individual subunit. That is, MCell repeatedly evaluates model rules against a given subunit’s flags (and those of the neighboring subunits) to determine which state transitions a subunit undertakes at each time step. In the following sub-sections, we describe all CaMKII model flags, the state transitions that apply to each flag, the conditions and rate parameters for each state transition, and related model assumptions. In Fig 2, we visually convey how CaMKII subunits transition between states according to our model’s rules. In S1 Appendix we summarize the state transition rules and rate parameter values.

**Fig 2.**
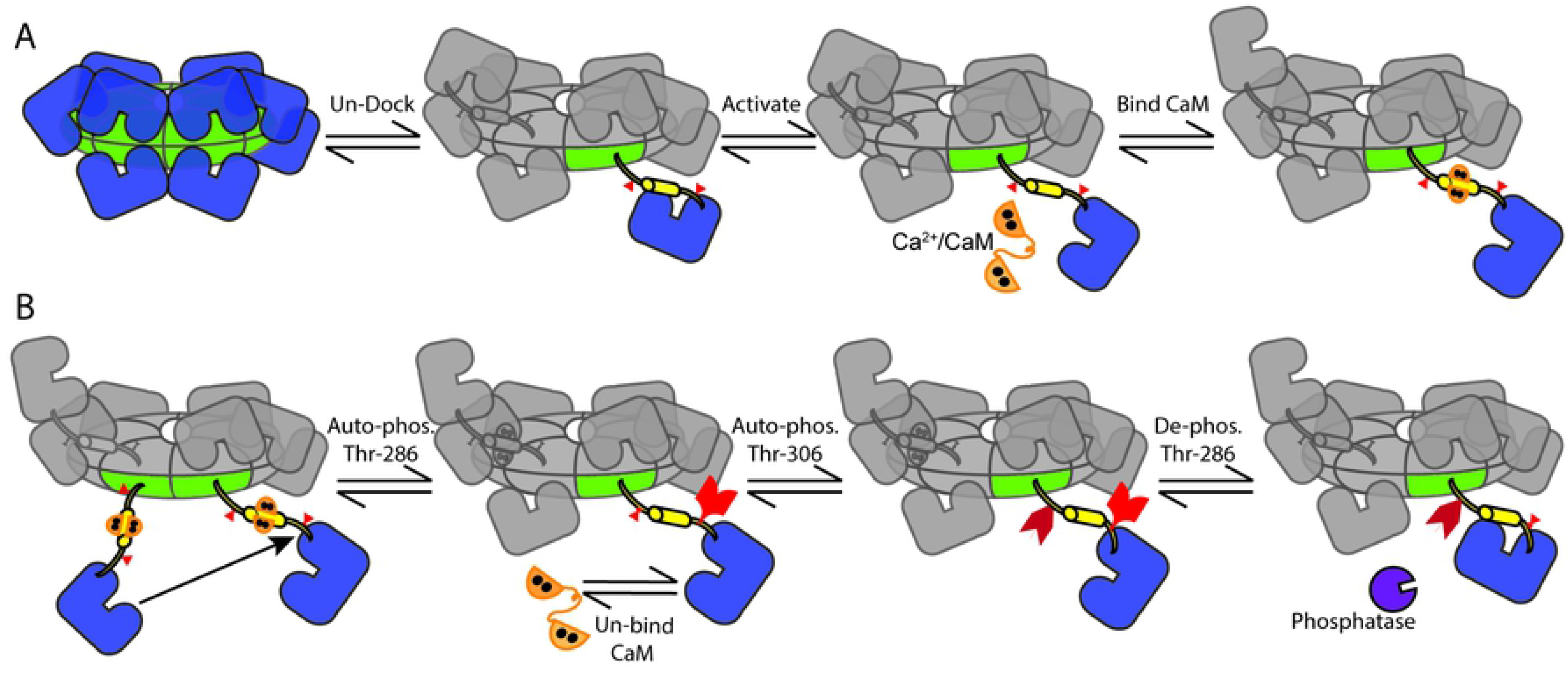
CaMKII holoenzyme state transitions. (A) CaMKII has twelve subunits arranged in two radially symmetric, directly apposed rings. Subunits may spontaneously undock/extend from the central hub or dock/retract (if inactive). When undocked, subunits may spontaneously open/activate. (B) If two neighboring subunits are active, one may auto-phosphorylate the other at Thr-286. If auto-phosphorylated (pThr-286), a subunit may remain active even upon un-binding of CaM. A pThr-286 subunit un-bound to CaM may additionally phosphorylate at Thr-306, blocking subsequent re-binding of Ca^2+^/CaM. A pThr-286 subunit may also bind and become de-phosphorylated by PP (purple).

#### Flag 1: Subunit docking

Docking is a binary flag that describes subunits as either “docked” or “undocked” to the CaMKII central hub. Subunits are instantiated in a docked state but may undergo numerous transitions between docked and undocked over the course of a simulation. At each time step, we assess a rule governing the subunit’s transition from a docked to undocked state. If this rule is satisfied, meaning that the subunit’s docking flag is verified as “docked”, then the transition is considered. Similarly, we assess a separate rule governing a transition from an undocked to docked state, which requires that the subunit not be bound to CaM and not phosphorylated at Thr-306 [17].

Subunit docking follows the structural model of Chao *et al*., who showed that a subunit cannot bind CaM as long as the subunit is in a compact conformation, docked to its central hub [17]. Docking implies a two-step process in which the subunit must first un-dock before subsequent CaM-binding, which accounts for the reported difference in binding rate for CaM to CaMKII-derived peptide (1 × 10^8^ M^−1^s^−1^ [50]) and for CaM to full-length CaMKII-T286A (1.8 × 10^6^ M^−1^s^−1^ [51]). Taking the ratio of these two rates gives an equilibrium constant for docking of 0.018, which is consistent with estimates by Chao *et al*., who assumed K_docking_ to fall between 0.01 and 100 [17]. With this equilibrium constant, we estimate kinetic rates for docking and undocking. For this estimation, we first note that subunit docking involves a structural conformation change on a relatively large scale. Referring to a separate, and notably smaller-scale, conformational change in our model, in which CaM quickly transitions from an initially- to fully-bound state (see Flag 3: CaM Binding), we assume the docked-to-undocked transition to proceed at an order of magnitude slower. We therefore arrive at an assumed rate for k_dock_ of 35 s^−1^. In turn, this gives an undocking rate k_undock_=k_dock_ × K_docking_ of 0.63 s^−1^, which lies within the range of 0.01 s^−1^ and 100 s^−1^ for k_undock_ assumed by Chao *et al*.

#### Flag 2: Subunit activation

The activation flag describes subunits as either “active” or “inactive”. An inactive subunit has no catalytic activity because the regulatory domain is bound to the subunit’s catalytic site; others may refer to it as a closed subunit. Conversely, an active subunit has catalytic activity because the regulatory domain’s inhibition of the kinase domain is disrupted; in other words, an active subunit is an open subunit. When a subunit is active, Ca^2+^/CaM and/or other proteins may access and bind CaMKII. In our model, the transition reaction from inactive to active (opening) involves no explicit rules (but rather occurs unconditionally and as governed by rates described below). In contrast, two rules inform the conditions for subunit inactivation: that the subunit is 1) not fully-bound to CaM, and 2) not phosphorylated at Thr-286.

To assign rate parameters for this flag, we first note that subunits can fluctuate between inactive and active states rapidly in the absence of Ca^2+^/CaM (on the order of hundreds of nanoseconds) [19, 52]. Noting this, we set the rate parameter for subunit inactivation at 1 × 10^7^ S^−^. Further, Stefan *et al*. determined that the activation probability (in the absence of CaM and phosphorylation) is 0.002, leading us to set our activation rate parameter to 2 × 10^4^ s^−1^ [29]. Thus, we arrive at a model in which CaMKII subunit activation is unstable until stabilized by CaM-binding or autophosphorylation.

#### Flag 3: CaM binding

CaM binding is a ternary flag meaning that each CaMKII subunit displays one of three states, where CaM may be “unbound”, “initially-bound”, or “fully bound”. Our model adapts previous work by Stefan *et al*. (2012) to describe CaM-binding to CaMKII as a two-step process [29]. First, CaM binds to the regulatory domain of a CaMKII subunit (residues 298-312), resulting in a low-affinity “initially bound” CaMKII state, which is compatible with both the closed and open subunit conformation. Second, if the initially bound CaMKII opens it may transition to a “fully bound” state that describes the complete, higher-affinity interaction between CaM and CaMKII along residues 291-312 (see Figure 5 in [29]). We specify three rules to govern the transition from an unbound to initially bound state: the subunit must be 1) undocked, 2) not PP-bound, and 3) un-phosphorylated at Thr-306. The transition reaction from initially bound to a fully bound state is governed by a single rule that the subunit already be active/open. Dissociation of CaM from a fully bound CaM-CaMKII state proceeds through the initially bound state before becoming completely unbound from CaMKII.

In order to determine the parameters governing initial binding of CaM to CaMKII, we use data on CaM binding to CaMKII-derived peptides, rather than full-length CaMKII. This is done to separate the intrinsic binding constants from the parameters governing subunit activation/inactivation and docking/undocking. The microscopic k_on_ for CaM binding to CaMKII has been measured, using a CaMKII peptide and fluorescently labeled DA-CaM, as 1 × 10^8^ M^−1^s^−1^ [50]. For the K_D_ governing initial CaM binding, we use the K_D_ reported by Tse *et al*. for CaM binding to a low-affinity peptide (CaMKII residues 300-312), which is 5.9 × 10^−6^ M [53]. From these two parameters, we can compute the dissociation rate of initially-bound CaM from CaMKII: k_off_CaM_ini_= K_d_CaM_ini_ × k_on_CaM_ = 590 s^−1^.

In order to determine the parameters governing the transition from initially-bound to fully-bound CaM to CaMKII, we note that this transition involves a structural compaction of the CaM molecule, which has been measured using fluorescent labels [50, 51]. Using fluorescent labels to analyze the structural compaction of CaM is convenient in its exclusion of effects due to conformational changes within CaMKII subunits or the CaMKII holoenzyme. Thus, we use these measurements as a proxy for CaM binding to a CaMKII peptide and to estimate parameters governing the transition between initially- and fully-bound CaM-CaMKII. Based on experiments by Torok *et al*., we identify a transition rate from initially- to fully-bound CaM-CaMKII of 350 s^−1^ and from fully- back to initially-bound CaM of 4 × 10^−3^ [50]. This means that, in the absence of obstructions to binding, the likelihood of a bound CaM molecule being in the initial binding state (rather than the fully bound state) is 4 × 10^−3^ / 350 = ∼1.1 × 10^−5^. This is consistent with a probability of CaM being bound to the high-affinity site of 0.99999 which was derived by Stefan *et al*. (2012) [29].

#### Flag 4: Phosphorylation at Thr-286

Phosphorylation at the residue Thr-286 is a ternary flag that describes this site as either “un-phosphorylated (uThr-286)”, “phosphorylated (pThr-286)”, or “phosphatase-bound”. We specify three rules to govern the reaction that transitions a subunit from uThr-286 to pThr-286: the subunit 1) be uThr-286, 2) be active, and 3) have an active neighbor subunit in the same holoenzyme ring. The neighboring subunit’s activation flag is considered because autophosphorylation is facilitated by its catalytic site. Our model only considers the counter-clockwise neighbor subunit because, in the absence of experimental observations to the contrary, we assume that steric effects cause autophosphorylation to occur in only one direction about a CaMKII ring, similar to previous work [54, 55]. The rate of autophosphorylation, 1 s^−1^, at Thr-286 is taken from an earlier study of CaMKII autophosphorylation in the presence of CaM [44].

De-phosphorylation of pThr-286 is facilitated by binding and enzymatic activity of protein phosphatases PP1 and PP2A, here referred to generally as PP [41, 42]. Two rules govern PP binding to a CaMKII subunit (the transition from pThr-286 to a phosphatase-bound state): that the subunit be 1) pThr-286 and 2) un-bound to CaM. It has been shown that a majority of autophosphorylated CaMKII in the PSD is dephosphorylated by PP1 [56, 57]; while in brain extracts autophosphorylated CaMKII is mostly dephosphorylated by PP2A [41]. The requirement that CaM be unbound from CaMKII in order for PP to bind to CaMKII is motivated by the observation that simultaneous binding of CaM and PP to the CaMKII regulatory domain may be mutually exclusive due to steric hindrance. CaM, having molecular weight 18 kDa, binds to the CaMKII regulatory domain around residues 290–309 [54, 58, 59], which is at least 4 residues, and at most 23 residues away from Thr-286 (again, see also Figure 5 in [29]). To the best of our knowledge, the peptide binding footprint of neither PP (PP1 nor PP2A) onto CaMKII is not yet fully described. However, both PP1 and PP2A are widely known to target pThr-286 [56, 57, 60] and de-phosphorylate threonine residues nearby alpha helices in other substrates [61, 62]. Additionally, the catalytic subunit of PP1 has a molecular weight of 37 kDa, which is nearly twice that of CaM and more than half that of a CaMKII subunit. Taken together, we hypothesize that the PP binding footprint likely overlaps with the CaM binding site, such that the presence of bound PP likely structurally excludes or impedes upon a subsequent binding of CaM to CaMKII. Similarly, the presence of bound Ca^2+^/CaM structurally would exclude coincident binding of PP. In S1 Appendix, we further discuss the quantitative basis of this structural exclusion hypothesis in light of the crystal structure of the PP1-spinophilin interaction (PDB: 3EGG) [63]. In short, PP1 tends to bind substrates at a site >20Å from the PP1 active site. Thus, if the PP1 binding footprint does not actually contain T286, then the furthest likely CaMKII residue of PP1 binding (at least on the hub domain side of T286) is G301, well within the CaM binding footprint (S1 Appendix). We examine the regulatory implications of this hypothesis by relaxing the rules of PP binding and requiring only that the subunit be pThr-286. The association, dissociation, and catalytic rates of PP for CaMKII are taken from Zhabotinsky (2000), using a Michaelis constant of 6 μM and a catalytic rate of 2 s^−1^[47].

#### Flag 5: Phosphorylation at Thr-306

Phosphorylation at the residue Thr-306 is a binary flag that describes this site as either un-phosphorylated (“uThr-306”) or phosphorylated (“pThr-306”). We model the transition from uThr-306 to pThr-306 using three rules: that that the subunit be 1) uThr-306, 2) active, and 3) un-bound by CaM. Our model uses a forward rate parameter 50-fold slower than that of phosphorylation at Thr-286, based on past experimental measurements [33, 64]. Over the course of our simulation times, we observe very few pThr-306 transitions and therefore exclude the reverse transition reaction describing de-phosphorylation of pThr-306 into uThr-306.

#### Stimulation frequency correlates with subunit activity

To validate our model, we assessed a variety of model outputs under various regimes of Ca^2+^/CaM stimulation. As a first assessment, we simulated a persistent Ca^2+^/CaM bolus, similar to experiments by Bradshaw *et al*. (2002), who monitored CaMKII autophosphorylation over time [55]. In Fig 3 we simultaneously monitored the time-course concentration of CaMKII subunit flags indicating: initially-bound Ca^2+^/CaM, fully-bound Ca^2+^/CaM, active CaMKII, and pThr-286. In the persistent, continuous presence of Ca^2+^/CaM, the concentration of subunits with initially-bound Ca^2+^/CaM (orange trace) is noisy and consistently low, implying that Ca^2+^/CaM transiently binds subunits in an initially-bound conformation. That is, initially-bound Ca^2+^/CaM seems rapidly to either dissociate or proceed to a fully-bound conformation. Fully-bound Ca^2+^/CaM (red trace) subunit concentrations closely follow those of active CaMKII subunits (dark blue trace) over time, providing evidence that Ca^2+^/CaM stabilizes CaMKII activation. Indeed, because the difference in concentrations of fully-bound Ca^2+^/CaM and active CaMKII is always small, we observe that although unbound CaMKII may spontaneously activate, these activated subunits rapidly return to an inactive state and are not likely to progress to a phosphorylated (pThr-286) state. We next observe that the increase of CaMKII autophosphorylation at Thr-286 (cyan trace) over time is strongly associated with increases in the number of subunits that are fully-bound to Ca^2+^/CaM and active subunits (dark blue and red traces, respectively). This is consistent with previous work showing that Ca^2+^/CaM must be bound to CaMKII for pThr-286 to occur [54] and CaMKII Ca^2+^-independent activity is strongly associated CaMKII autophosphorylation at Thr-286 [17, 51, 65, 66]. Furthermore, we observe in Fig 3A that more than 80 percent of CaMKII subunits are autophosphorylated at t=20sec, which is of similar magnitude and timescale as observed by Bradshaw *et al*. (see Figure 2A in [55]).

**Fig 3.**
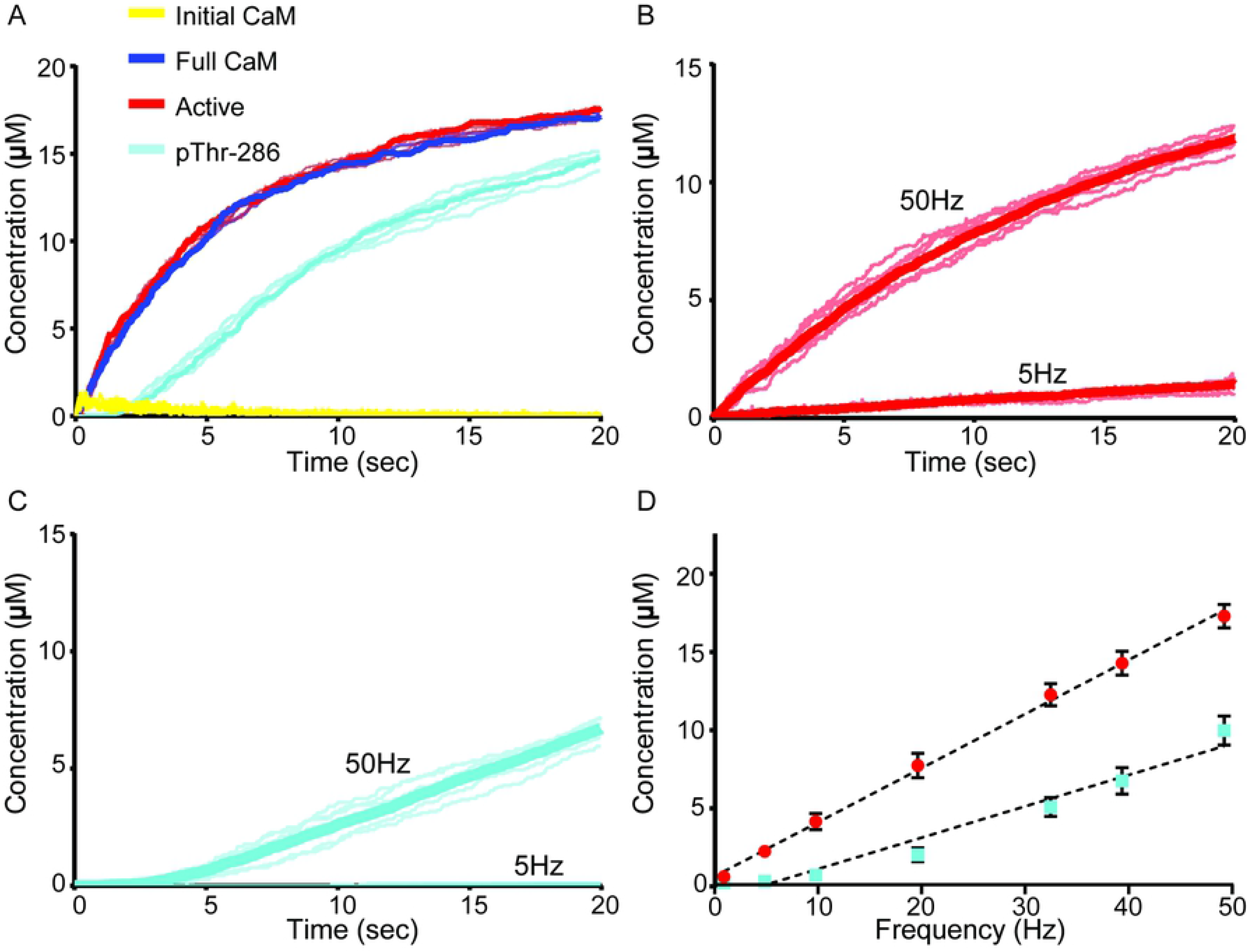
Validation of the Rule-based Model. Bold traces (A-C) and solid circles (D) are the average of N = 50 executions. For each species (A-C), six representative traces are also shown (semi-transparent lines). (A) Model output resulting from stimulation with a large continuous bolus of Ca^2+^/CaM. Concentrations of active (red), initially CaM-bound (yellow), fully CaM-bound (blue), and pThr-286 (cyan) subunits. (B) Time-course average concentration (bold trace) of active subunits stimulated by 5 Hz or 50 Hz Ca^2+^/CaM. (C) Time-course concentration of pThr-286 subunits stimulated continuously by 5 Hz or 50 Hz Ca^2+^/CaM. (D) Frequency-dependent activation (red) and pThr-286 (cyan) of CaMKII subunits, with SEM error bars. Black dotted traces are linear fits.

Next, we assessed model behavior under low- and high-frequency stimulating conditions. CaMKII activation and autophosphorylation at Thr-286 in response to 5Hz and 50Hz Ca^2+^/CaM is plotted in Figure 3B and 3C, respectively; 50 seeds were run for each condition, with 6 representative traces (transparent lines) and the average response (bold) plotted. As expected, the data showed significantly greater levels of CaMKII activation and autophosphorylation at 50Hz [12, 20]. Indeed, we compared our result in Fig 3C to work by Shifman *et al.* (2006), who observed much lower autophosphorylation at low Ca^2+^/CaM concentrations (less than 2 μM) than at high concentrations (see Figure 4D in [44]). Therefore, because our 50Hz model cumulatively exposes CaMKII to approximately ten times as much Ca^2+^/CaM per second as our 5Hz model, our results in Fig 3C are consistent with Shifman *et al*., showing much higher autophosphorylation at 50Hz than 5Hz.

**Fig 4.**
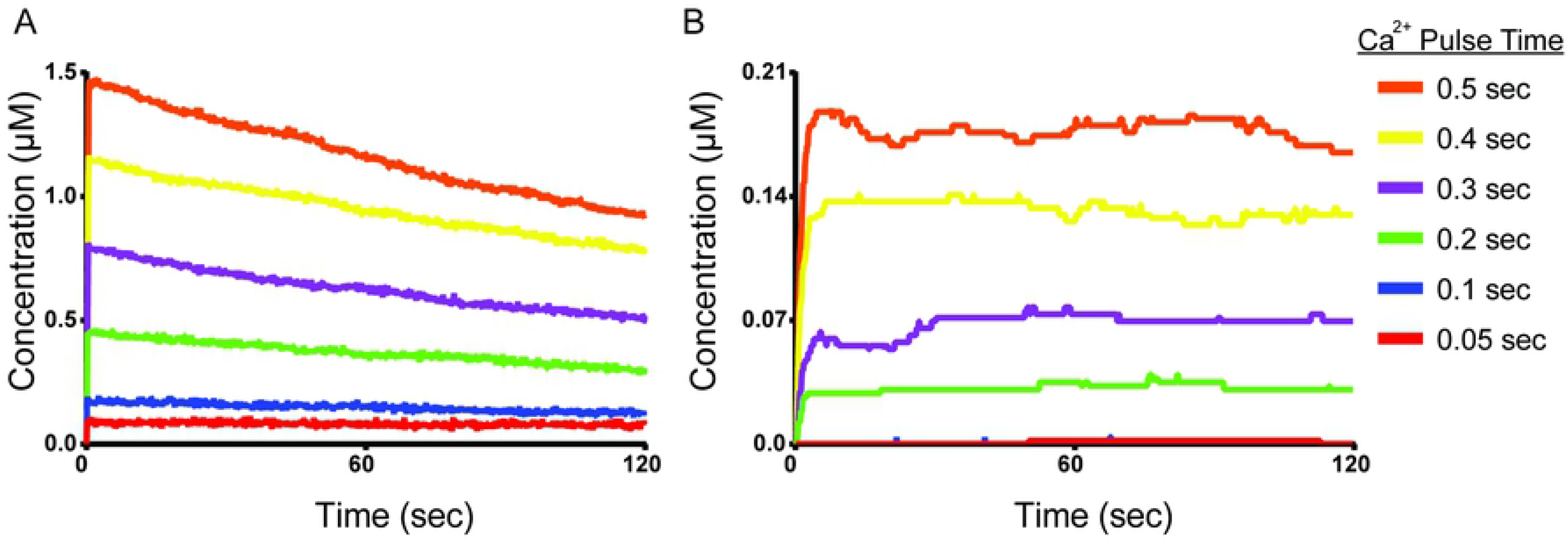
Response to short Ca^2+^/CaM pulse stimulation. Average concentration of (A) active and (B) pThr-286 CaMKII subunits over time, following Ca^2+^/CaM stimulating pulses of length 0.05 (red), 0.1 (blue), 0.2 (green), 0.3 (purple), 0.4 (yellow), and 0.5 (orange) seconds. Each trace represents the average of N=50 executions. See Appendix S1 for identical data shown with SEM error bars and over the first two seconds of simulated time.

To further determine how stimulation frequency affects CaMKII activity, the model was stimulated at frequencies ranging from 1Hz to 50 Hz. At each frequency, models were sampled at 20 seconds of simulation time. We observe a nearly linear correlation between both subunit activation (R^2^ = 0.99) and pThr-286 (R^2^ = 0.96) and stimulation frequency (Fig 3D). This result is consistent with computational results from Chao *et al*., who developed a stochastic model that also yielded a linear relationship between pThr-286 and stimulation frequency for frequencies greater than 1 Hz [15]. Taken together, these results (Fig 3) show that our model behaves as expected and is able to produce CaMKII activity and autophosphorylation behaviors similar to previous computational and experimental results.

#### Exploring Switch-like Behavior in CaMKII

CaMKII has long been theorized to exhibit switch-like or bistable behavior, which could underlie the importance of pThr-286 to learning and memory formation [4, 47, 48, 67, 68]. However, experimental efforts have struggled to identify a bistability between CaMKII and phosphatase activity. Though recently, Urakubo *et al*. used the chelator EGTA to control single pulses of Ca^2+^ in a mixture of CaM, CaMKII, PP1, and NMDAR peptides, leading to what seemed to be the first direct observation of CaMKII bistability [69]. Referring to Urakubo *et al*., we explored whether a spatial stochastic model of the CaMKII dodecamer would exhibit near bistability or switch-like behavior for concentration parameters of Ca^2+^, CaM, CaMKII, and PP known to exist in hippocampal spines. To explore this bistability, we stimulated the model with a set of short Ca^2+^/CaM input pulses (which could also be reproducible *in vitro*). Importantly, we did not aim to identify true bistability because exploring the many combinations of Ca^2+^, kinase, and phosphatase concentrations was outside the scope of this paper. Instead we wondered if, by stimulating with brief pulses of Ca^2+^/CaM of variable duration, our model would exhibit switch-reminiscent pThr-286 behavior. Specifically, we predicted a Ca^2+^/CaM stimulation threshold below which pThr-286 was unachievable and above which pThr-286 was maintained.

In Fig 4 we exposed our model to single Ca^2+^/CaM pulses of constant magnitude but of variable duration (similar to Figure 1B in [69]). The model was stimulated with single Ca^2+^/CaM input pulses of magnitude 26 μM and varying duration (0.05, 0.1, 0.2, 0.3, 0.4, or 0.5 sec). Different pulse durations resulted in distinct levels of subunit activation, where longer pulse durations resulted in greater activation and autophosphorylation (p-Thr286) levels, (Fig 4A and B, respectively). Interestingly, subunits stimulated by even the shortest pulses of 0.05 or 0.1 sec, appeared to sustain their activation for the complete simulation period (120 sec). However, these short-pulse (0.05-0.1 sec) stimulations rarely resulted in autophosphorylation (pThr-286, Fig 4B). Longer (0.2-0.5 sec) Ca^2+^/CaM pulses resulted in greater levels of subunit activation that started declining immediately after the Ca^2+^/CaM pulse ended (Fig 4A), but elicited pThr-286 levels that were generally sustained for the duration of a simulation (Fig.4B). Taken together, we found that CaMKII may be thresholded at a level of Ca^2+^/CaM exposure below which pThr-286 is unobserved and above which pThr-286 is achieved and subsequently sustained across several minutes even in the presence of phosphatase.

We briefly explored how this Ca^2+^/CaM threshold may depend on the number of directions by which subunits can autophosphorylate their neighbors. Note that in the results up to this point, autophosphorylation was limited to occurring in a single direction, or degree of freedom. That is, subunits could only autophosphorylate their adjacent neighbors [17]. We therefore created alternative versions of our model in which autophosphorylation could occur with multiple degrees of freedom, both intra- and/or trans- holoenzyme ring. We used these higher-degree of freedom models to monitor the rates of pThr-286 formation both in bulk and on an individual subunit basis. As expected, pThr-286 formation and intra-holoenzyme propagation rates increased with increasing degrees of freedom (see S1 Appendix, Figure S2.2), though the differences would likely not be distinguishable by bench-top experimentation. In addition, the length of time in which consecutive subunits remained autophosphorylated also increased with increasing degrees of freedom. This implied that subunits may be more frequently autophosphorylated on time-average with increasing degrees of freedom (See Appendix S1, figures S2.3 and S2.4). Future experimental and computational studies could perhaps explore autophosphorylation with higher degrees of freedom.

Figure 4 suggested a threshold of Ca^2+^/CaM activation beyond which CaMKII remains autophosphorylated, implying a balance between kinase and phosphatase activity. We wondered how a putative balance between CaMKII autophosphorylation and phosphatase activity might be regulated. In the previous experimental work by Urakubo *et al*., maximally-phosphorylated CaMKII was maintained in the presence of PP1 and GluN2B peptide for as long as 8 hours (at 4°C). In that work, addition of the kinase inhibitor K252a to phosphorylated CaMKII resulted in a steady decline in pThr-286 towards basal levels, suggesting that maintenance of pThr-286 over time was not due to low phosphatase activity, but rather a recovery of de-phosphorylated subunits back a phosphorylated state. To recreate inhibition of kinase activity in our model, at time t=30 sec we introduced a high concentration (21 μM) of K252a, enough to bind all CaMKII subunits in the model. K252a binding results in a blocked CaMKII state that cannot be autophosphorylated (see Flag 2 in S1 Appendix). Importantly, the blocked CaMKII subunit can still be de-phosphorylated at pThr-286. In separate simulations we explored the effects of a phosphatase inhibitor, which was also introduced at t=30sec. To simulate the introduction of a phosphatase inhibitor, we defined the catalytic rate of de-phosphorylation by PP1 (k_cat_^PP1^) as a time-dependent variable that assumed a value of zero at t = 30sec. This implementation of kinase and phosphatase inhibition preserved normal CaM and PP1 binding dynamics.

We compared non-inhibited, control versions of our model to versions in which kinase activity or phosphatase activity was inhibited after stimulating with Ca^2+^/CaM for 2 sec (Fig 5). As expected, inhibiting phosphatase activity (green trace) caused kinase activity to dominate, resulting in a steady increase in pThr-286 compared to the control (black trace). Surprisingly, the kinase-inhibited model (blue trace) showed little difference compared to the control. Instead of causing phosphatase activity to dominate, simulated kinase inhibition caused pThr-286 activity to persist as if no kinase inhibitor were present. Because pThr-286 persisted even in the presence of kinase inhibitor, we hypothesized that some other, non-enzymatic mechanism in our model was contributing to the maintenance of pThr-286.

**Fig 5.**
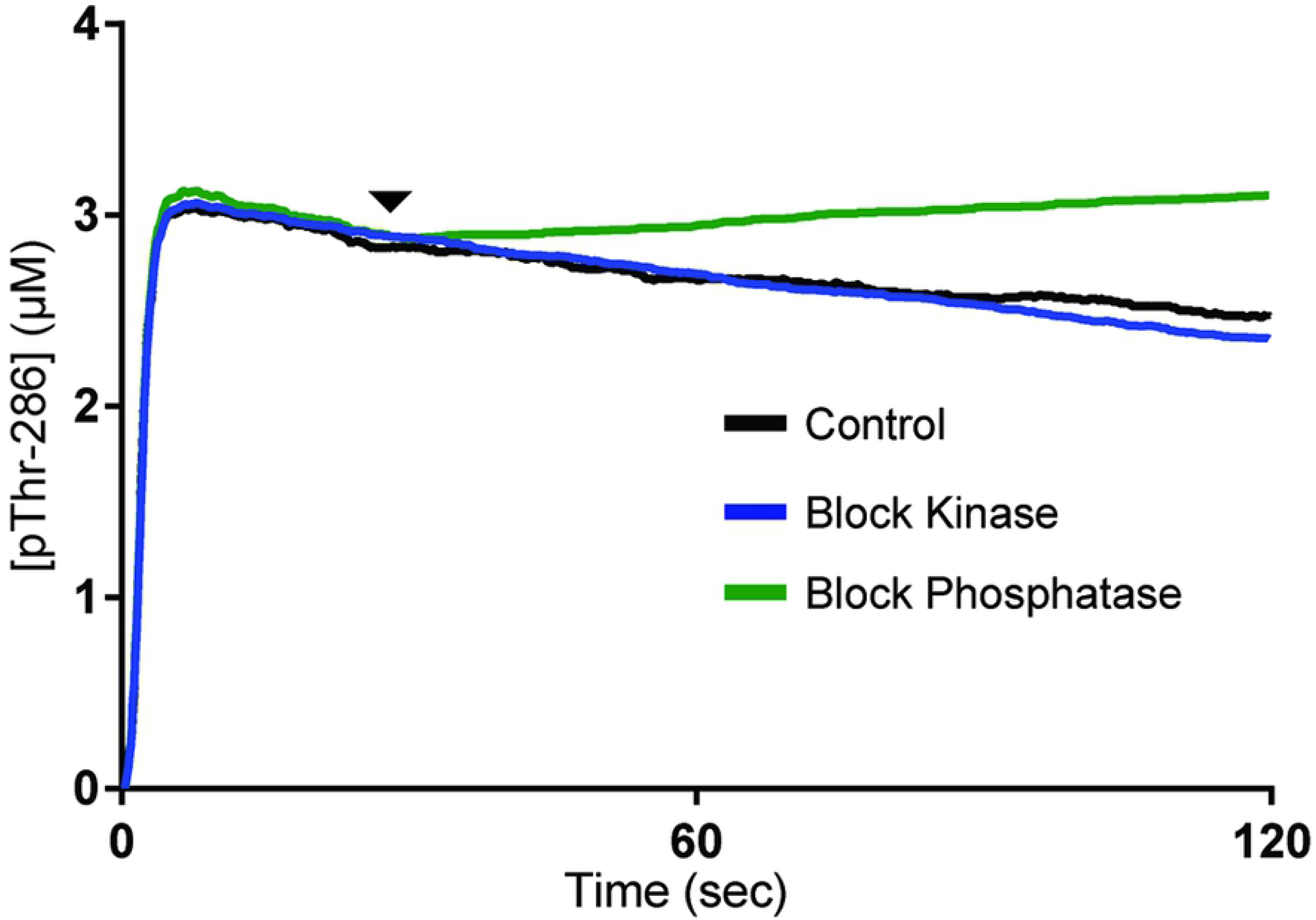
Blocking kinase or phosphatase activity. Average concentration of pThr-286 CaMKII subunits over time. For all traces, the model is stimulated by a 2 sec pulse of Ca^2+^/CaM. At time t=30 sec (arrowhead), either a kinase inhibitor (blue trace) or phosphatase inhibitor (green trace) is introduced. No inhibitor is introduced in the control (black trace). Each trace represents the average of N=50 executions.

### CaM-dependent exclusion of PP1 binding stabilizes autophosphorylation

To understand why our model as-presented in Fig 5 showed no significant response to kinase inhibition, we wondered if another mechanism was regulating the putative balance between kinase and phosphatase activity. In every simulation presented thus far, we assumed that CaM binding to the CaMKII regulatory domain sterically hinders PP binding to the regulatory domain, and vice-versa. This was implemented in the model via a rule that requires a subunit be unbound by CaM in order for PP to bind.

To test the role of PP exclusion by CaM, we created a second version of our model in which PP binding would become allowable regardless of the presence of CaM. In contrast to our original “exclusive” model, the “non-exclusive” model required only that a subunit be pThr-286 in order for PP binding to be allowable. In other words, the non-exclusive model allowed Ca^2+^/CaM and PP to bind CaMKII agnostically of each other. Aside from this rule adjustment, our exclusive and non-exclusive models utilized identical parameters. As in Fig 5, we selected a Ca^2+^/CaM bolus time of 2 sec. Again, we monitored both CaMKII activation (Figs 6A and 6B) and pThr-286 (Figs 6C and 6D) over 120 seconds of simulated time. Critically, both the exclusive and non-exclusive models were examined with high (purple trace) and low (orange trace) association rate parameter values for PP binding to CaMKII. Increasing and decreasing the association rate of PP (k_on_^PP^ is normally set to 3 μM^−1^sec^−1^) to CaMKII by one order of magnitude accounted for parameter uncertainty and provided a magnified view of the signaling effects of CaM-mediated exclusion of PP binding.

Our results suggested that CaM-dependent exclusion of PP is an important regulatory mechanism for maintaining CaMKII autophosphorylation levels. While the PP exclusion rule had little to no effect on CaMKII subunit activation (Fig 6A and Fig 6B), pThr-286 (Fig 6C and Fig 6D) was highly influenced by the PP exclusion rule. In the exclusive model (Fig 6C), pThr-286 levels were steady and stable despite varying the PP association rate parameter by two orders of magnitude. In contrast, the non-exclusive model (Fig 6D) showed that for a high PP association rate, significant pThr-286 levels were never achieved. Moreover, for a low PP association rate, the non-exclusive model briefly attained pThr-286 levels similar to those achieved in the exclusive model, but the pThr-286 levels then declined while also displaying a high level of noise. It seemed that in order to maintain pThr-286 over longer time periods, CaMKII required a mechanism regulating phosphatase access, and a regulator of phosphatase access could be CaM itself.

**Fig 6.**
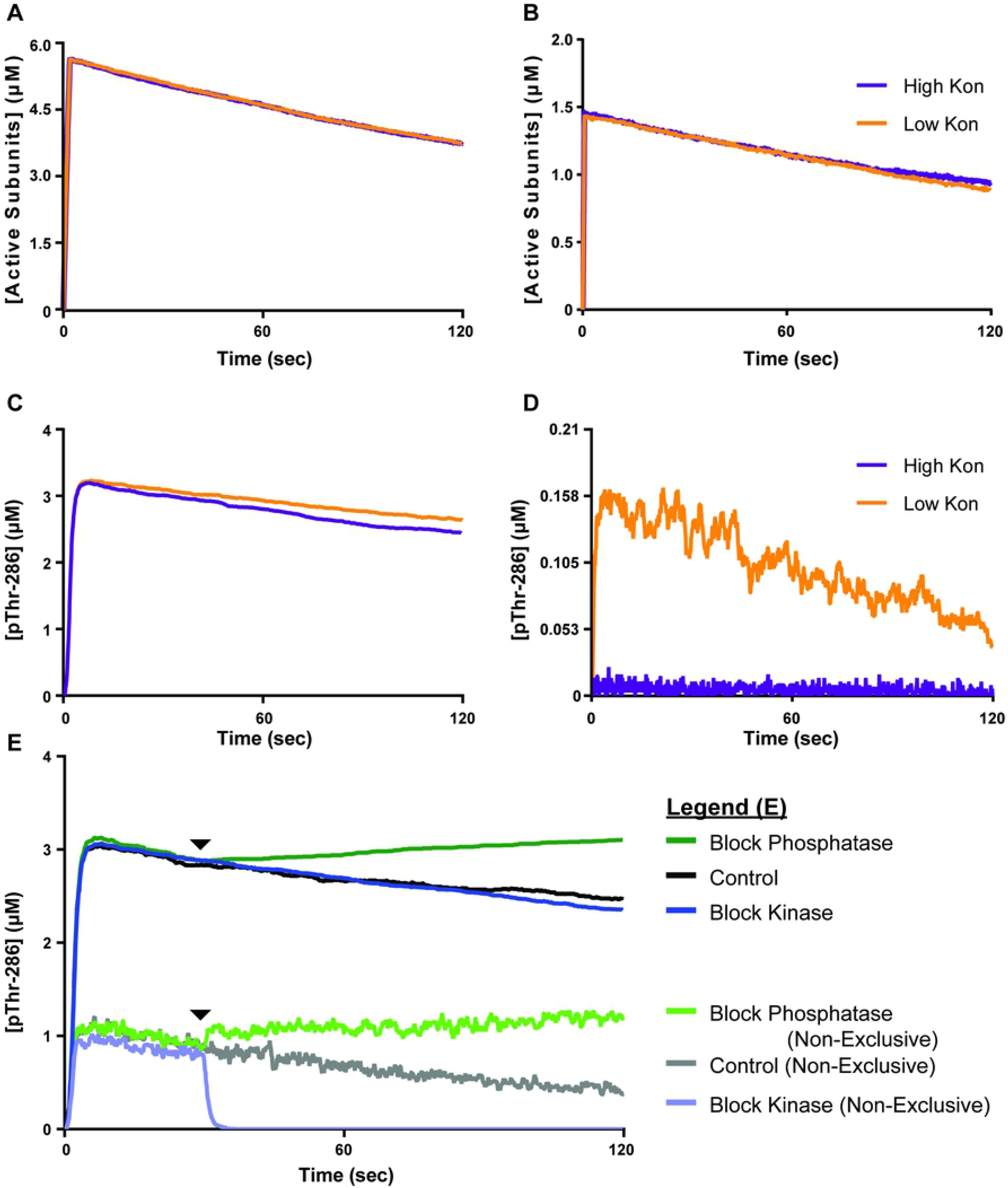
Comparison of Exclusive and Non-exclusive Models. For all traces, models are stimulated by a 2sec pulse of Ca^2+^/CaM. (A) Active CaMKII subunits over time in our exclusive model. (B) Active CaMKII subunits over time in our non-exclusive model. (C) pThr-286 subunits over time in our exclusive model. (D) pThr-286 subunits over time in our non-exclusive model. (A-D) The parameter value for the rate of PP association (k_on_^PP1^) with CaMKII is either increased (purple traces) or decreased (orange traces) by one order of magnitude. (E) Extension of Fig 5 to include non-exclusive model results. At time t=30sec (arrows), either a kinase inhibitor (light blue trace) or phosphatase inhibitor (light green trace) is introduced. No inhibitor is introduced in the control (grey trace). All traces are the average of N=50 executions.

To reinforce our assertion that CaM-dependent structural exclusion of PP binding stabilizes pThr-286, we repeated simulations shown in Fig 5, but with our non-exclusive model. In Fig 6E, we stimulated our non-exclusive model with a 2sec pulse of Ca^2+^/CaM and then monitored pThr-286 over time. For these simulations, k_on_^PP1^ was restored to its standard value of 3 μM^−1^sec^−1^. As in Fig 5, in separate simulations we inhibited at t=30sec either phosphatase activity, kinase activity, or neither (control). The control (grey trace) was reminiscent of results in Fig 6D, in which pThr-286 was achieved but then slowly declined on a steady yet noisy basis. Notably, all non-exclusive model variants were much noisier than their exclusive model counterparts in Fig 6E. Inhibiting phosphatase activity (light green trace) in the non-exclusive model again caused kinase activity to dominate and pThr-286 levels to generally increase over time, similarly to the exclusive model. In contrast to the exclusive model, inhibiting kinase activity (light blue trace) in the non-exclusive model rapidly and totally abolished pThr-286. It seemed that for the non-exclusive model, in which CaM and PP could bind simultaneously, inhibiting kinase activity caused phosphatase activity to dominate. Taken together, these results suggested that in addition to supporting CaMKII subunit activation, CaM also has a role in maintaining CaMKII activity by blocking phosphatase access and thereby slowing down dephosphorylation.

## Discussion

In this work, we use rule- and particle-based methods with the software MCell to model the complete CaMKII holoenzyme. Rule-based modeling allows us to account for and monitor multiple CaMKII states simultaneously without eliciting combinatorial explosion. By explicitly accounting for multiple CaMKII states, we can use this model to explore regulatory mechanisms such as the CaM-dependent maintenance of pThr-286 by structural exclusion of phosphatase binding to CaMKII.

Previous multi-state models of CaMKII exist but are different in focus and in scope from the present model. For example, our model is based on an earlier multi-state model by Stefan *et al*. (2012) [29] implemented in the particle-based stochastic simulator StochSim [70]. StochSim accounts for subunit topology (i.e. the user can specify whether a subunit is adjacent to another, and reactions can be neighbour-sensitive), but StochSim does not explicitly account for spatial information. MCell, as a spatial simulator, offers more possibilities to precisely account for spatial effects and to situate models in spatially realistic representations of cellular compartments. In addition, the model by Stefan *et al*. provides only for interactions between adjacent CaMKII molecules on the same hexamer ring and therefore models CaMKII as a hexamer, not a dodecamer. Similarly, another previous model of CaMKII by Michalski and Loew (2012) uses the softwares BioNetGen and VCell to offer an infinite subunit holoenzyme approximation (ISHA) of the CaMKII hexamer [71–73]. The ISHA asserts that under certain enzymatic assumptions, the output of a multi-state CaMKII model is independent of holoenzyme size when the number of subunits exceeds six. However, Michalski’s ISHA model is most suitable for systems containing only one holoenzyme structure-dependent reaction such as the autophosphorylation at Thr-286. Additional reactions to describe actin binding [74] or subunit exchange [14, 15] may invalidate Michalski’s ISHA, whereas our model can in the future readily accommodate additional, holoenzyme structure-dependent phenomena. Finally, a more recent rule-based model of the CaMKII holoenzyme by Li and Holmes [26] offers a detailed representation of how CaM binds to Ca^2+^ and subsequently activates CaMKII subunits, based on earlier results of CaM regulation [75]. Li and Holmes offer valuable and detailed insight into how CaM binding to CaMKII depends on Ca^2+^ dynamics. While our model is less detailed in representing the regulation of CaM itself, our model is much more detailed in representing other aspects of CaMKII regulation, including multiple modes of CaM binding, conformational change, detailed holoenzyme structure, multiple phosphorylation sites, and dephosphorylation. We can in the future expand our MCell model to account for multiple holoenzyme structure-dependent phenomena and simultaneously incorporate this model into the broader Ca^2+^-dependent signaling network.

This work in-part demonstrates the value of MCell as a rule-based modeling framework. Rule-based modeling accommodates much larger state spaces than is possible using conventional systems of differential equations. Admittedly, not all models (including models of CaMKII) require extensive state spaces, but rule-based modeling results can help justify the assumptions typically used to reduce a state space. For example, our model conditions yield, as shown in Fig 3A, negligible levels of initially-bound CaM compared to other states such as fully-bound CaM or pThr-286. Therefore, it might sometimes be appropriate to exclude an initially-bound CaM state from future implementations in frameworks for which combinatorial explosion is a concern. Aside from addressing combinatorial explosion, rule-based models are especially well-suited to discern otherwise concealed mechanisms, as exemplified by Di Camillo *et al*. who used rule-based models to identify a robustness-lending negative feedback mechanism in the insulin signaling pathway [49]. Furthermore, MCell describes CaMKII holoenzymes as discrete particles in space, which will lend realism to future spatial-stochastic models of Ca^2+^-dependent signaling networks in the dendritic spine, a compartment in which the Law of Mass Action is invalid [24]. This particle-based framework also allows for individual subunit monitoring, which works in conjunction with the Blender software plugin, CellBlender (see S1 Movie).

One of the results of this work is the identification of distinct levels of CaMKII activation and pThr-286 in response to distinct pulses of Ca^2+^/CaM stimulation. Distinct levels of CaMKII activation could tune the selectivity of CaMKII for certain downstream binding targets such as AMPA receptors or the structural protein PSD-95. If stimulation-dependent tuning of CaMKII activation were observed, it would be reminiscent of other studies that have implicated feedback loops [27] and binding dynamics [24] as regulators of Ca^2+^-dependent enzyme activation. For example, a recent study suggests that competition is an emergent property that tunes the Ca^2+^ frequency dependence of CaM binding to downstream targets, leading Ca^2+^/CaM to set distinct levels of calcineurin- and CaMKII-binding [12]. Similarly, CaMKII itself could preferentially select downstream binding partners as a function of its level of activation by Ca^2+^/CaM, possibly providing a mechanism by which CaMKII facilitates certain LTP-related molecular events. Additionally, our observation of distinct levels of CaMKII activation and thresholded pThr-286 could be an indication of long-hypothesized switch-like behavior in synaptic plasticity [4, 67]. If switch-like behavior in fact occurs, then pThr-286 is likely maintained by a balance in kinase and phosphatase activity.

While investigating a putative interplay in CaMKII kinase and PP phosphatase activity in maintaining pThr-286 levels, we may have identified a CaM-dependent mechanism that blocks PP binding to CaMKII. In a model that excludes simultaneous binding of CaM and PP to CaMKII, pThr-286 significantly increases upon phosphatase inhibition, yet in the same model kinase inhibition causes little change in pThr-286 over time (Fig 5). In contrast, a non-exclusive model that allows simultaneous binding of CaM and PP shows that introduction of a kinase inhibitor rapidly abolishes pThr-286. These results suggest that CaM-dependent exclusion of PP may provide a stabilizing mechanism. Additionally, we use our MCell-based implementation of the model to monitor transitions between multiple states of distinct subunits within holoenzymes (Fig 7 and S1 Movie).

**Fig 7.**
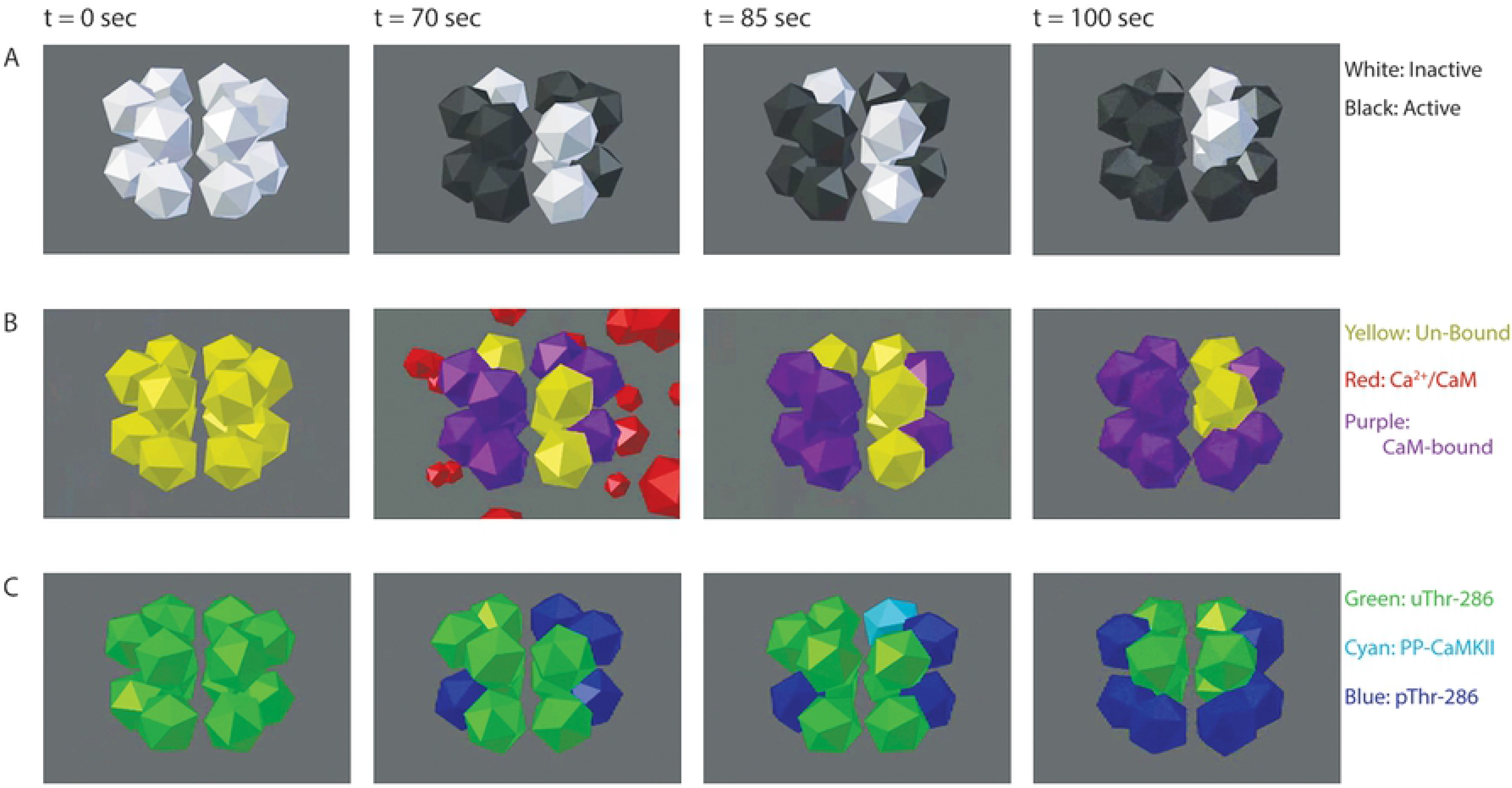
Visualizing Individual Subunits with MCell and CellBlender. In the exclusive model, PP does not bind a pThr-286 subunit until Ca^2+^/CaM dissociation (see t = 85 sec, comparing rows B and C). Each frame depicts the same CaMKII holoenzyme, from the same perspective, at identical time points under 50Hz Ca^2+^/CaM stimulation. Each dodecahedron is a single CaMKII subunit. (A) Inactive CaMKII subunits (white) spontaneously become active (black) and remain active while bound to Ca^2+^/CaM. (B) Un-bound CaMKII subunits (yellow) will not bind Ca^2+^/CaM (red) and become Ca^2+^/CaM-bound (purple) unless the subunit had previously activated. (C) uThr-286 subunits (green) become pThr-286 (blue). If Ca^2+^/CaM dissociates from a pThr-286 subunit, then PP can bind and form a PP-CaMKII complex (cyan).

The major outcome of this work is a proposed mechanism in which bound Ca^2+^/CaM could exclude PP from accessing CaMKII subunits, thereby protecting pThr-286. We assert that CaM-dependent exclusion of PP could provide a functional role for so-called “CaM trapping” [54] and possibly contribute to CaMKII bistability. Indeed, a model by Zhabotinsky (2000) explored CaMKII bistability, indicating that two stable states of pThr-286 would in-part require very high CaMKII concentrations, seemingly to bolster kinase activity in the system [47]. However, the Zhabotinsky model assumes that CaM and PP1 could bind CaMKII simultaneously, possibly exaggerating the ability of PP1 to de-phosphorylate at Thr-286. If PP1 binding were to be encumbered in the Zhabotinsky model, perhaps through CaM-dependent exclusion, then bistability might be achievable at lower CaMKII concentrations.

Previous studies have sought to explore the dependence of CaMKII de-phosphorylation on the presence of Ca^2+^/CaM. An experiment by Bradshaw *et al*. (2003) quantifies PP1-mediated de-phosphorylation rates of pThr-286 *in vitro*, in the presence or absence of the Ca^2+^ chelator EGTA (see Figure 4B in [76]). The Bradshaw results suggest that PP1 activity at 0°C is unaffected by the presence of bound-CaM to CaMKII, seemingly at odds with the results of our model. However, EGTA can only act on free Ca^2+^, Ca^2+^/CaM has a low dissociation rate for Ca^2+^, and phospho-CaMKII has a low dissociation rate for Ca^2+^/CaM. Also, other studies [12, 24, 44] indicate that sub-saturated Ca^2+^/CaM may in fact significantly bind CaMKII. Thus, because the Bradshaw experiment does not remove CaM from the reaction mixture, the Bradshaw results may be confounded by a slow unbinding of CaM from CaMKII. Our results motivate a revised phosphatase assay, which a) uses physiological temperatures, b) ensures detection of de-phosphorylation solely of pThr-286, and c) totally removes CaM from the reaction mixture.

Future work may need to explore the ability of PP1 to bind CaMKII in the presence of sub-saturated Ca^2+^/CaM. Referring to Fig 6, although some CaM may remain bound to CaMKII when Ca^2+^ concentration decreases, the resulting sub-saturated (lower affinity) Ca^2+^/CaM state might be out-competed by PP1. In other words, if only the CaM C-terminus were bound to CaMKII, would PP1 have sufficient structural access and/or affinity to bind also? Thus, in addition to the revised phosphatase assay mentioned above, further structural studies of the CaM-CaMKII and PP1-CaMKII interaction are needed. Also, because this work could only model multi-state CaMKII (but not also multi-state CaM) due to limitations in MCell 3.3, perhaps a future version of MCell should provide for the diffusion of multiple multi-state proteins. With a platform that can handle multiple multi-state proteins, a model could much more explicitly handle Ca^2+^/CaM-binding and further explore our results.

CaM-dependent PP exclusion could provide an added layer of robustness to similar mechanisms that may protect pThr-286 from de-phosphorylation. For example, Mullasseril *et al*. (2007) observe that endogenous, PSD-resident PP1 cannot de-phosphorylate CaMKII at pThr-286, whereas adding exogenous PP1 does cause de-phosphorylation [68]. The results by Mullasseril *et al*. suggest that endogenous PP1 is somehow sequestered by the PSD scaffold, and only upon saturation of this scaffold by exogenous PP1 does pThr-286 become de-phosphorylated. Our results indicate that perhaps in addition to saturating the PSD scaffold, the added exogenous PP1 could be out-competing CaM for binding to CaMKII, thereby terminating protection of pThr-286 by CaM. As another example, Urakubo *et al*. suggest that pThr-286 could be protected from PP activity by GluN2B binding, showing that GluN2B peptides are necessary for an apparent CaMKII bistability *in vitro* [69]. Notably, Urakubo *et al*. observe a decline in pThr-286 upon kinase inhibition, in contrast with our exclusive model, though this is likely due to differences in the conditions and timescales between Urakubo’s experiments and our model. Overall, it seems scaffold-dependent sequestration of PP1 [68], GluN2B-dependent PP exclusion [69], and CaM-dependent PP exclusion could together provide considerable robustness of pThr-286 to phosphatase activity.

## Methods

### Simulation Methods

In each MCell execution, proteins are instantiated at time zero having random positions within a 0.32 μm^3^ (0.32 fL) cube. All proteins are described as three-dimensional volume molecules having the following concentrations: 1.52 μM CaMKII (30 holoenzymes, 360 subunits), 22.8 μM CaM (450 discrete proteins), and 0.86 μM PP1 (17 discrete proteins). Because CaMKII particles are modeled using the specialized COMPLEX_MOLECULE syntax and MCell 3.3 does not accommodate diffusion for such particles, CaMKII is given no diffusion constant. In contrast, CaM and PP1 are simple volume-type molecules that move about the model space with a diffusion constant 6 × 10^−6^ μm^2^/sec. All models are run at a time step of 0.1 μs for a total of either 20 or 120 seconds of simulation time, depending on the model variant. For statistical significance, all model variants are repeated 50 times each.

CaM activation/inactivation is modeled by a pair of forcing functions which serve as a proxy for Ca^2+^ flux. Both forcing functions are time-dependent square waves and inform the rates at which free CaM transitions between states. Equation 1 rapidly transitions all free CaM towards an active (Ca^2+^/CaM) state, and Equation 2 rapidly transitions all free CaM towards an inactive (apo-CaM) state.

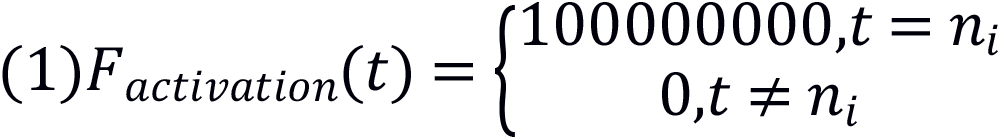

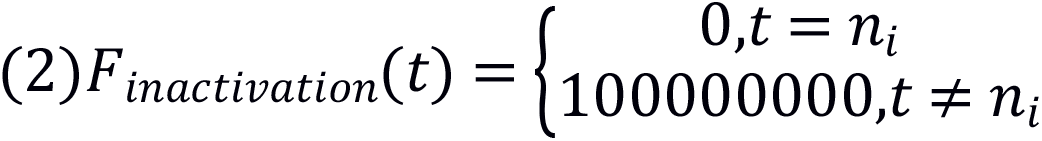

For both Equations 1 and 2, n = i/f where i is the number of time step iterations and f is frequency. Time t iterates at 0.01sec intervals for the complete duration of a simulation. Equations 1 and 2 therefore yield a peak width of 0.01sec regardless of frequency, which allows us to directly compare the effect of different Ca^2+^/CaM frequencies on CaMKII activity, without having to account for variable amounts of Ca^2+^/CaM exposure per pulse. In separate simulations without frequency dependence (i.e. Ca^2+^/CaM is continuously available to CaMKII), Equation 1 is adjusted to always fulfill the t=n_i_ condition. Similarly, for pulse simulations in which Ca^2+^/CaM becomes withdrawn or blocked, Equations 1 and 2 are given abbreviated time domains.

All MCell code and associated files are available online at Github, the Purdue University Research Repository (DOI: 10.4231/MBPK-D277), and the University of Edinburgh Repository.

## Acknowledgements

The authors offer special thanks to Neal Patel for many helpful conversations and assistance creating the CaMKII movie visualizations in CellBlender. We also thank Stefan Mihalas, Nicolas LeNovere, Elizabeth Phillips, Kaisa Ejendal, David Umulis, and Tyler VanDyk for their helpful advice and comments on the manuscript.

## Supporting Information

### S1 Appendix. Appendix

This document enumerates the model parameters, discusses combinatorial explosion, shows alternative visualizations of select data, and discusses the quantitative basis for PP1/CaM mutual exclusion from CaMKII binding.

